# KinAce: a web portal for exploring kinase-substrate interactions

**DOI:** 10.1101/2023.12.08.570875

**Authors:** John A. P. Sekar, Yan Chak Li, Avner Schlessinger, Gaurav Pandey

**Affiliations:** Department of Genetics and Genomic Sciences, Icahn Genomics Institute, Icahn School of Medicine at Mount Sinai, New York, NY 10029, USA; Department of Pharmacological Sciences, Icahn School of Medicine at Mount Sinai, New York, NY 10029, USA; Department of Artificial Intelligence and Human Health, Icahn School of Medicine at Mount Sinai, New York, New York 10029, USA

**Keywords:** network visualization, knowledge discovery, data sharing, post-translational modification databases, cell signaling, systems biology

## Abstract

Interactions between protein kinases and their substrates are critical for the modulation of complex signaling pathways. Currently, there is a large amount of information available about kinases and their substrates in disparate public databases. However, these data are difficult to interpret in the context of cellular systems, which can be facilitated by examining interactions among multiple proteins at once, such as the network of interactions that constitute a signaling pathway. We present KinAce, a user-friendly web portal that integrates and shares information about kinase-substrate interactions from multiple databases of post-translational modifications. KinAce enables the visual exploration of these interactions in systems contexts, such as pathways, domain families, and custom protein set inputs, in an interactive fashion. We expect KinAce to be useful as a knowledge discovery tool for kinase-substrate interactions, and the aggregated KinAce dataset to be useful for protein kinase studies and systems-level analyses. The portal is available at https://kinace.kinametrix.com/.

## Introduction

Kinase proteins and their substrates are critical to cell signaling^1–4^. Dysregulation of kinases is involved in multiple pathologies, such as cancer, neurodegeneration, cardiovascular disease and infectious diseases, making kinases and their substrates priority targets for drug development^5–7^. Kinase-substrate interactions are known to be complex, where, for example, a particular kinase can phosphorylate multiple substrates, or a particular substrate can be phosphorylated by multiple kinases at one or more site(s)^4,8,9^. Understanding kinase function requires examining these interactions at multiple scales, from their structural aspects^10–13^, to their participation in specific pathways^14–18^, to the systems-level effects that they produce^3,4,9,19^.

Currently, there is a wealth of information available about kinase-substrate interactions disseminated by a number of public tools and web portals. Some focus on molecular features of kinases, such as their structural conformations (e.g., KinaMetrix^10^ and KinCore^11^) and druggability (e.g., KLIFS^12^). Others focus on molecular features of substrates targeted by kinases, specifically phosphorylation sites (e.g., PhosphoSitePlus^20^, iPTMNet^21^ and EPSD^22^) and motifs (e.g., The Kinase Library^13^). There are also portals that focus on phylogenetic classification of kinases (e.g., KinHub^23^ and Coral^24^) and information about understudied kinases (e.g., The Dark Kinome Knowledgebase^25^). Additionally, extensive information about kinase interactions is presented in protein databases like UniProt^26^ and BioGRID^27^. However, only a relatively small fraction of the available data on these interactions is interpretable in a functional context. For example, the vast majority of experimentally known phosphorylation sites have no associated kinase(s)^20,21^. Of the known kinase-substrate interactions, only a few well-understood ones have been incorporated into curated pathway databases (e.g., KEGG^28^, Reactome^29^ and PathwayCommons^30^). Thus, there is an opportunity to uncover new kinase functionality by examining the aggregated molecular-level data on these interactions from a systems perspective. Visualizing and analyzing kinase-substrate interactions in the context of functional groupings like pathways can yield useful knowledge about kinase biology that can be leveraged for applications like drug discovery.

To address this important need, we have developed the KinAce web portal for aggregating, sharing and visualizing kinase-substrate interactions in the human genome. KinAce aggregates and shares a comprehensive dataset of kinase-substrate interactions from PhosphoSitePlus^20^, iPTMNet^21^ and EPSD^22^, which are three large databases of post-translational modifications with recent and regular updates, and, which also provide coverage of several other data sources. KinAce provides multiple ways to visualize these interactions in varied functional/systems contexts, such as pathways, domain families and custom sets of genes provided by the user (e.g., from gene expression studies). Collectively, the data and visualization capabilities provided by KinAce represent a unique resource for exploring kinase-substrate interactions that complement the current ecosystem of tools for analyzing kinase data.

## Materials and Methods

### Aggregating kinase-substrate interaction data

The set of known human kinase-substrate interactions is continuously evolving. As a consequence, there are a large number of kinase-substrate interaction databases that overlap with each other significantly^31^. They are also maintained to different extents, and are standardized, e.g., in the proteins names they use, to different extents.

To build a comprehensive dataset of kinase-substrate interactions, we selected resources capturing the largest amount of public information, and with the most recent and regular updates. Additionally, it was important that the resources we selected contain references for each interaction, typically to the original publication, for data provenance. Three resources fit our criteria:

1. PhosphoSitePlus^20^, the largest continuously maintained database of expert-curated kinase-substrate interactions;
2. iPTMNet^21^, a database that curates information from PhosphoSitePlus and PhosphoELM^32^, as well as information extracted by text-mining of scientific literature;
3. EPSD^22^, an annotated collection of multiple curated databases, including PhosphoELM, PSEA^33^, PostMOD^34^, and RegPhos^35^, as well as a subset of PhosphoSitePlus.

We aggregated kinase-substrate interactions from the most current versions of the respective databases as of October 4^th^, 2023 (PhosphoSitePlus v6.7.1.1, iPTMNet v5.0, EPSD v1.0). To standardize protein and gene names, we cross-referenced them with protein identifiers from the reviewed subset of the UniProt human proteome^26^ and gene symbols from HGNC^36^ current on the same date. Additionally, we incorporated kinase-specific data from KinHub^23^, Coral^24^ and the Dark Kinome Knowledgebase^25^, such as memberships of kinase proteins in phylogenetic groups^37^.

Each entry in the aggregated KinAce dataset is a triplet involving a kinase, a substrate and the site on the latter phosphorylated by the kinase. From these, we extracted unique kinase-substrate pairs, which in turn constitute the kinase-substrate interaction network used for visualizations on KinAce. The full interaction dataset, along with associated information like the source database(s), can be downloaded from the portal directly.

### The KinAce portal and interface

KinAce is a web-based portal for sharing and visualizing the network of human kinase-substrate interactions (Figure 1). The portal is available at https://kinace.kinametrix.com/.

**Figure 1:**
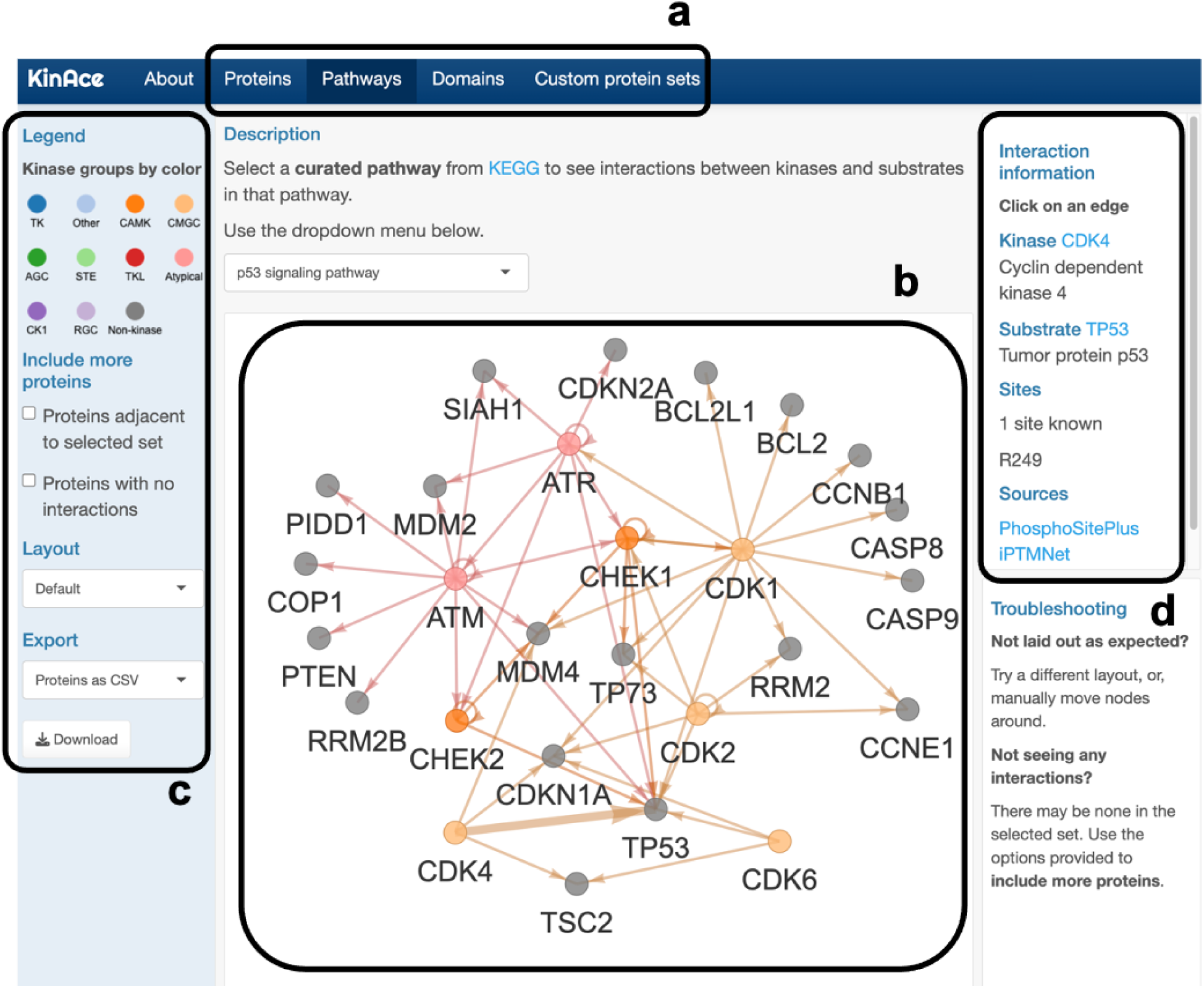
KinAce functionalities. The KinAce web portal aggregates, shares and visualizes kinase-substrate interactions in the human genome from established databases of post-translational modifications. **(a) Choosing sets of proteins**. The tabs highlighted provide different ways to select sets of proteins for which kinase-substrate interactions will be displayed: individual proteins and their interactors, curated pathways from KEGG, protein families mapped to InterPro domains, and custom protein sets provided by the user. **(b) Visualizing kinase-substrate interactions.** Interactions within the selected set are shown as a network of directed edges from kinase nodes to substrate nodes. Kinase nodes are colored by kinase group and non-kinase nodes are colored gray. **(c) Interacting with the visualization.** The left sidebar shows the legend for node colors, an option for including proteins that are one degree away from the selected set, an option for unhiding nodes without edges, layout and export options. **(d) Provenance of data.** When the user selects an edge in the displayed network, the right panel displays information about the corresponding interaction, including links to the database sources from which the interaction was obtained. The user can follow these links to retrieve the literature reference(s) that reported that interaction. The specific example shown in this figure is the p53 signaling pathway visualized in the Pathway tab with the CDK4-TP53 interaction selected.

One of the main features of KinAce is the ability to visualize known kinase-substrate interactions within selected sets of proteins as a network diagram. KinAce provides multiple ways to select these sets of proteins (Figure 1a), organized into four main tabs on the portal:

1. select a single protein on the Proteins tab,
2. select one of several curated pathway from KEGG^28^ on the Pathways tab,
3. select one of several InterPro^38^ domains on the Domains tab, and
4. provide a set of proteins on the Custom protein sets tab.

The network diagram produced (Figure 1b) displays kinase and non-kinase proteins in the selected set as colored and gray nodes respectively. Kinase nodes are colored by their phylogenetic group membership^23,24,37^. Interactions from kinases to their respective substrates are represented as directed edges. The user can move nodes around, as well as zoom in or out, to find the most meaningful layout for the displayed network. The left sidebar (Figure 1c) includes additional layout choices, as well as the option to unhide proteins with no interaction data. The user can also expand the network to include additional proteins adjacent to those in the selected set. Finally, the visualized network can be downloaded as an image (PNG), table of interactions (CSV) or in commonly used graph formats (GML^39^, GraphML^40^ and DOT^41^).

Notably, data can be traced back to their original source(s). When the user clicks on an edge in a network diagram, the panel on the right (Figure 1c) displays information about the corresponding interaction, including

- the name of the kinase, and a link to its UniProt page,
- the name of the substrate, and a link to its UniProt page,
- a list of the known sites on the substrate phosphorylated by the kinase, and,
- links to the original database source(s) from which the interaction was obtained.

The user can follow these links to retrieve the primary literature reference(s) supporting each interaction. In fact, we used the KinAce web portal to retrieve all supporting references for specific kinase-substrate interactions mentioned in this paper.

## Results and Discussion

### The KinAce dataset

The data aggregated from the most recent versions of PhosphoSite, iPTMNet and EPSD consisted of 16,360 unique kinase-substrate-site triples, representing 8,365 unique kinase-substrate pairs involving 416 kinases and 2,707 non-kinases. When we analyzed the original data source of each pair, we found that the three databases overlapped substantially, but also contributed several unique interactions individually (Figure 2a). The majority of interactions in the resulting dataset were between kinases and non-kinases, but there were also a notable number of autophosphorylations (n=229) and interactions between non-unique pairs of kinases (n=1,016) (Figure 2b). A small number of kinases and substrates dominated a large number of interactions (Figure S1). The full dataset can be downloaded directly from the portal.

**Figure 2:**
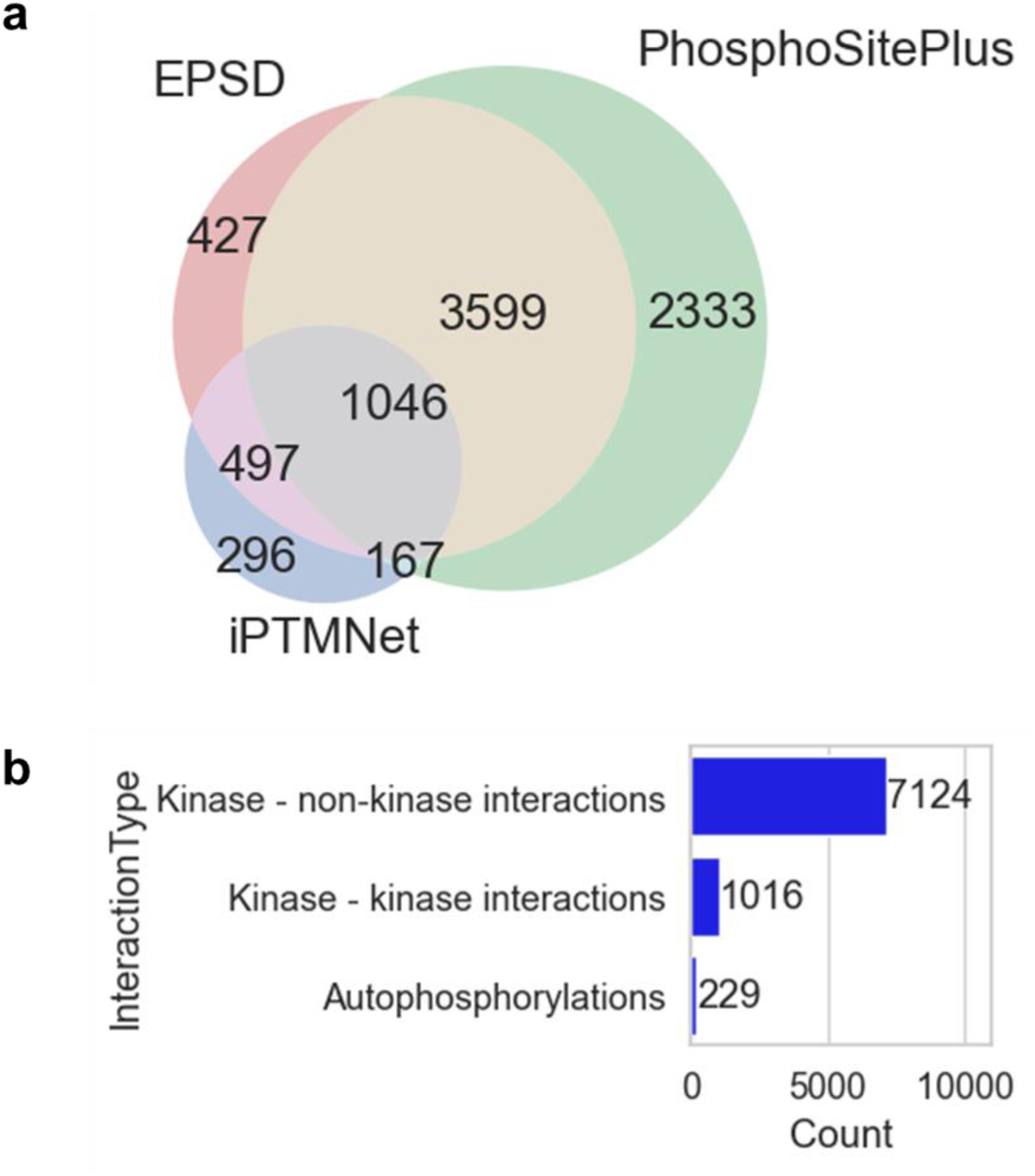
Kinase-substrate interaction statistics in the KinAce dataset. KinAce aggregated a dataset of 8,365 kinase-substrate interactions from the PhosphoSitePlus, EPSD and iPTMNet databases. **(a) Breakdown of interactions by data sources.** The three sources overlapped, but also contributed unique interactions to the KinAce dataset. **(b) Breakdown of data by interaction types**. The majority of the interactions include interactions between kinases and non-kinases (n=7,124), but a substantial number of interactions either involved two different kinases (n=1,016), or were autophosphorylations (n=229).

### Exploring interactions of individual proteins with KinAce

KinAce can be used to identify kinase-substrate interactions among sets of proteins of interest to the user.

The Proteins tab displays the known interactions of a protein selected by the user, which may be a kinase or a non-kinase. The network produced is a system-wide summary of kinase regulation involving the protein. For example, consider the visualization centered on BRAF, a well-known oncoprotein and component of the MAPK signaling pathway^42^ (Figure 3a). Outgoing edges from BRAF highlight substrates of BRAF, which not only include the canonical MAP2K1 and MAP2K2 kinases, but also non-kinases with diverse functions, such as the apoptosis promoter BAD^43^, the translation elongation factors EEF1A and EEF1A2^44^, and the ion transporter SLC9A1^45^. Incoming edges to BRAF indicate kinases that regulate BRAF, such as AKT1^46^ and AKT3^47^, as well as feedback from downstream proteins like MAPK1 and MAPK3^48^. This visualization is also effective for non-kinases. For example, the network centered on SLC9A1 (Figure 3b) displays the diverse range of kinases it interacts with in addition to BRAF, such as RPS6KA1 and ROCK kinases^49^, as well as AKT1^50^ and MAPK1^51^. Finally, the visualization permits traversing the network: double-clicking on a node recenters the visualization on the corresponding protein.

**Figure 3:**
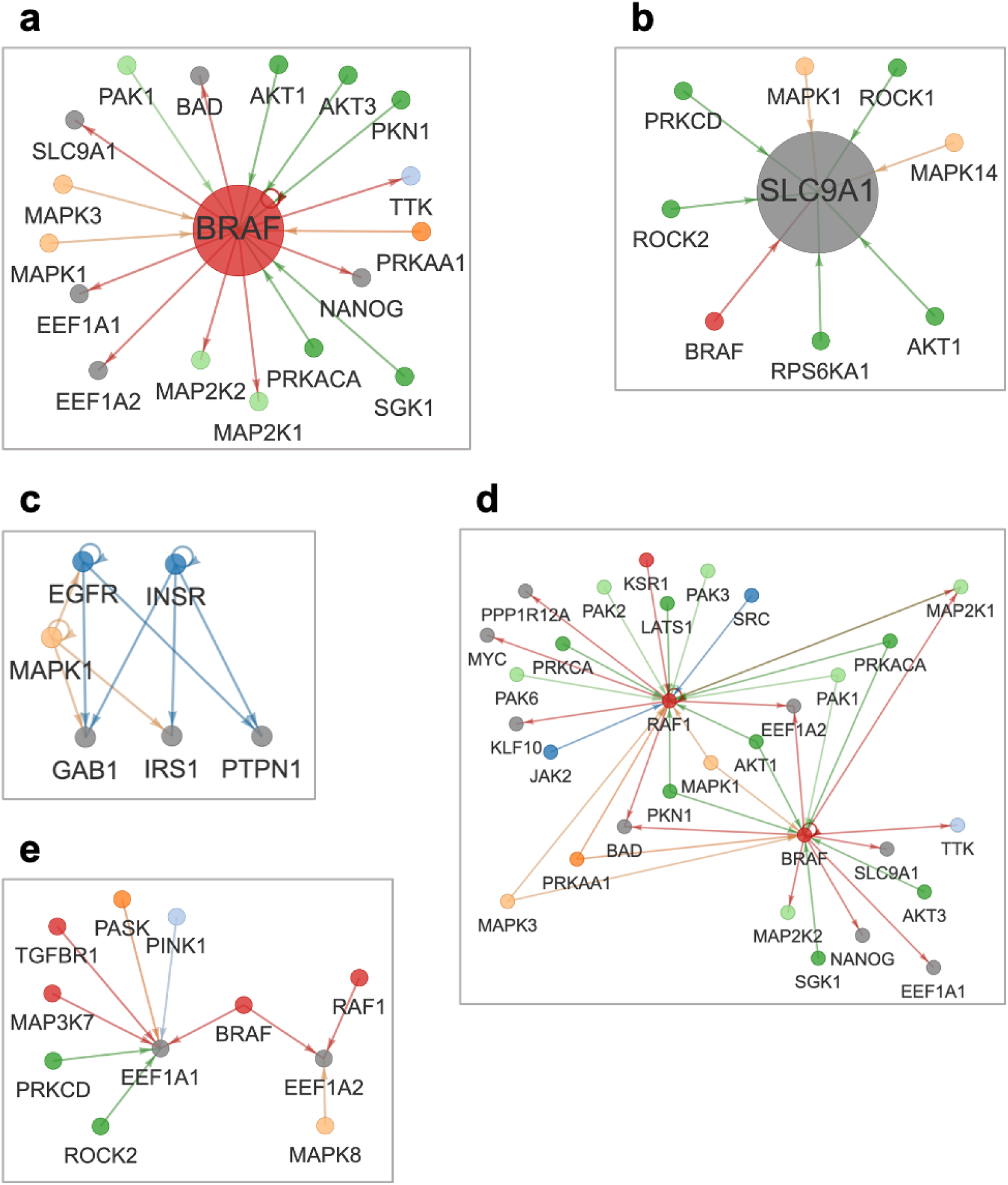
Exploring kinase-substrate interactions of proteins using KinAce. The Proteins tab displays all interactions of a selected protein. **(a) Interactions of the kinase BRAF**. Outgoing edges point to proteins phosphorylated by BRAF, such as MAP2Ks and non-kinases EEF1As and SLC9A1. Incoming edges indicate kinases that phosphorylate BRAF such as AKTs as well as downstream MAPKs. **(b) Interactions of the non-kinase SLC9A1.** Incoming edges indicate interactions with multiple kinases such as BRAF, AKT1 and ROCK1. The Custom protein sets tab lets the user specify any set of proteins within which to visualize interactions. The user can expand the network by selecting the option to include proteins adjacent to the selected set. **(c) EGFR-INSR crosstalk.** A custom set of proteins focused on overlap between EGFR and INSR pathways. **(d) Comparing RAF kinases**. The expanded network for the custom set of BRAF and RAF1 can be used to elucidate their shared versus exclusive interactions. **(e) Comparing EEF1A proteins**. The expanded network for the custom set of EEF1A1 and EEF1A2 shows the differences in the interactions they have with various kinases.

The Custom protein sets tab visualizes the kinase-substrate interactions among a set of proteins provided by the user. This allows the user to generate networks for any functional context of interest. For example, a user interested in the overlap between insulin receptor (INSR) and EGFR pathways^52^ can display the interactions involving the receptors EGFR and INSR and a select set of downstream proteins like the kinase MAPK1, the scaffold protein GAB1, insulin substrate IRS1 and the phosphatase PTPN1 (Figure 3c).

To explore additional interactions relevant to the proteins of interest, the user can select the option in the sidebar to include kinases and substrates that interact with the proteins shown in a network. For example, when we enable this option for the custom set of BRAF and RAF1, the network produced includes several new proteins (Figure 3d). Notably, it enables a systems-level comparison of BRAF and RAF1: we can delineate interactions common to BRAF and RAF1 on the diagram (e.g., MAP2K1) versus interactions unique to them (e.g., AKT3 and SRC). This approach is useful for non-kinases also. For example, we can see that the dataset includes more interactions with kinases for EEF1A1 than for EEF1A2 (Figure 3e).

### Exploring pathways with KinAce

KinAce can be used to examine kinase-substrate interactions among proteins in specific pathways of interest, as well as make inferences.

Specifically, the Pathways tab enables visualizing and analyzing kinase-substrate interactions in several curated pathways from the KEGG database^28^. For example, when examining the network produced for the p53 signaling pathway (Figure 1), one can identify various kinases involved in checkpoint signaling^53^ such as checkpoint kinases (CHEK1 and CHEK2), DNA damage sensors (ATM and ATR), multiple cyclin-dependent kinases (CDKs) and others. Additionally, one can identify several important non-kinases including tumor suppressors TP53 and TP73, the oncogene MDM2, caspases (CASP8, CASP9), cyclins (CCNB1, CCNE1), among others. This view of the pathway is complementary to other visualizations^28–30^ used by the community, as it focuses on kinase-substrate interactions.

The grid layout option can be effective for exploring pathways with large protein sets, such as MAPK signaling (Figure S2). However, for several pathways, the defined protein sets include many kinase substrates but not kinases. In these cases, the sidebar option to include adjacent proteins can be useful for discovering regulatory interactions. For example, expanding the currently disconnected KinAce network of the folate biosynthesis pathway (Figure 4a) revealed kinases that target metabolic enzymes in the pathway (Figure 4b), such as CAMK2 kinases targeting SPR^54^, an enzyme involved in several disease pathologies^55^.

**Figure 4:**
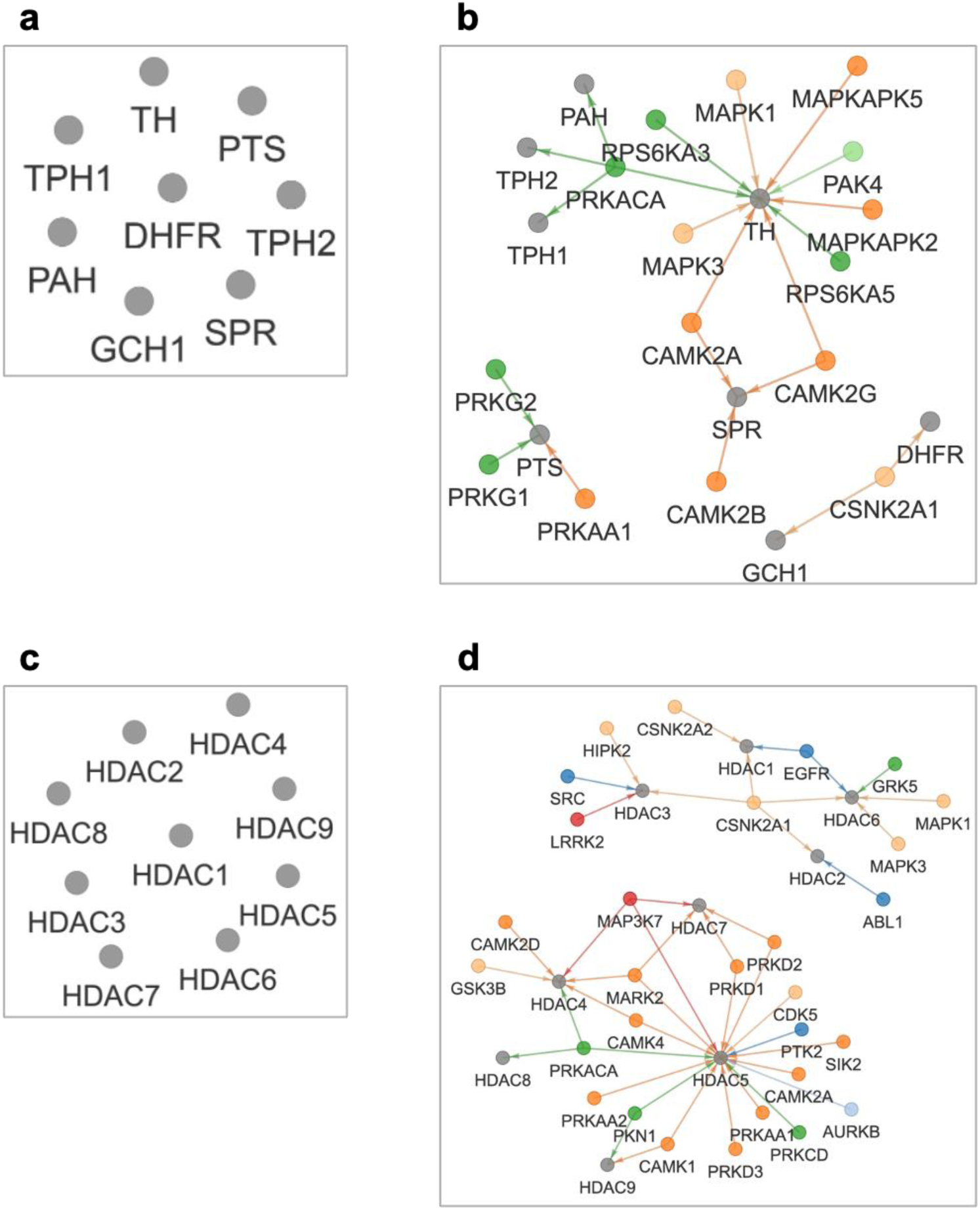
Exploring kinase-substrate interactions in functional contexts, such as pathways and shared domains using KinAce. The Pathways tab displays interactions from a selected pathway from KEGG database. The Domains tab displays interactions of proteins that share a selected domain. If the selected protein sets do not include kinases, expanding the network to include adjacent proteins can reveal regulatory interactions. **(a) The folate biosynthesis pathway**. This network, selected in the Pathways tab, shows a number of metabolic enzymes and no kinase-substrate interactions. **(b)** The **expanded view** reveals regulatory interactions such as phosphorylation of SPR by CAMK2 kinases. **(c) Proteins with the histone deacetylase domain.** This network, selected in the Domains tab contains multiple HDAC proteins and no kinases. **(d)** The **expanded view** reveals the wide range of kinases that phosphorylate HDACs, as well as other interactions.

Finally, to assess the general usefulness of the KinAce data for understanding pathways, we examined the enrichment of KEGG pathways in the kinase substrates included in the dataset using the Enrichr platform^56,57^. This analysis highlighted 201 significantly enriched pathways in 6 KEGG categories (Figure S3a), including environmental information processing (receptor-and small-molecule-activated pathways), genetic information processing (transcription, replication, DNA repair etc.), metabolism, cell-scale phenomena (cell cycle, autophagy, motility etc.), organism-level systems (endocrine, digestive, circulatory etc.), and a range of human pathologies, such as cancer, infectious disease, cardiovascular disease and neurodegeneration. The full set of enriched pathways and the corresponding p-values are reported in Appendix 1.

The above examples and results demonstrate the utility of KinAce and its underlying data for systems biology, specifically pathway analyses.

### Exploring domain groups with KinAce

KinAce can be also used to examine domain composition of kinases and substrates from a systems perspective.

Specifically, the Domains tab enables selecting sets of kinases and substrates that share InterPro domains^38^, and potentially highlight shared patterns of regulation. For example, the network for the SH3 domain (Figure S4) highlights its close association with Src-type kinases like SRC, FYN, FGR, LCK, LYN etc^58^. Additionally, InterPro domains can serve as an entry point for examining kinase regulation in a functional context. For example, phosphorylation of histone deacetylases (HDACs) is essential to normal physiological function^59^, and in recent years, was found to be disrupted by pathogens like SARS-CoV2^60^. Selecting the option to include adjacent proteins expands the currently disconnected KinAce network of the histone deacetylase domain (Figure 4c) and enables a broader examination of kinases that phosphorylate HDACs (Figure 4d).

Finally, to assess the general utility of KinAce and its data for functional studies based on domain composition, we conducted an enrichment analysis of the kinase substrates included therein against InterPro domain annotations using Enrichr^56,57^. The analysis produced 147 significantly enriched InterPro terms. Although there is no universally accepted functional classification of domain terms, we identified five general functional categories among the enriched terms (Figure S3b): nuclear functions (e.g., DNA-binding, RNA recognition and other mechanisms involved in transcription, replication, chromatin remodeling, etc.), cytoplasmic proteins with primarily protein-binding function (e.g., adaptors, scaffolds, small-molecule sensors, and ligand receptors), proteins with catalytic function (e.g., kinases, phosphatases, phosphodiesterases, lipid kinases, GTPases, and peptidases), structural proteins that are part of the cytoskeleton and extracellular matrix, and membrane proteins with transport function (e.g., ion channels and permeases). The full set of enriched domain terms and the corresponding p-values are reported in Appendix 2.

The above examples and results highlight the usefulness of the KinAce dataset for users interested in protein function represented by domain composition.

## Conclusion

The KinAce web portal is a user-friendly resource for exploration and systems analysis of interactions between human protein kinases and their substrates. KinAce aggregates and shares kinase-substrate interactions from several established databases of post-translational modifications, and helps visualize this dataset as interactive networks. Specifically, the portal provides multiple ways to specify protein sets of interest for visualization, such as interactomes of individual proteins, proteins organized into pathways, proteins sharing domains, and user-defined custom protein sets. Individual interactions highlighted on these visualizations can be traced back to database sources and their corresponding literature references. The aggregated KinAce dataset is a useful resource for future kinase studies and systems-level analyses, as was demonstrated using functional enrichment analysis of these data with biological pathways and domains. The results from the dataset and the visualization features highlighted in this paper illustrate the utility of the KinAce resource for systems-level study and applications of kinases and their substrates.

## Acknowledgements

We thank David Stein for assisting with web portal deployment, as well as Noah Herrington for proof-reading this manuscript and providing useful comments. This work was supported by NIH grant U01CA271318.

## Author Contributions

Conceptualization, Data Curation, Writing – Original Draft, J.A.P.S.; Visualization, Software, J.A.P.S. and Y.C.L.; Writing – Review & Editing, J.A.P.S., A.S., and G.P.; Resources, Supervision, Project Administration, Funding Acquisition, A.S., and G.P.

## STAR Methods

### Resource Availability

#### Lead contact

For more information, please contact Gaurav Pandey at.

#### Data and code availability

The KinAce portal is available at https://kinace.kinametrix.com/. The paper aggregates and analyzes publicly available data. The aggregated dataset and open-source code for the portal is maintained at https://github.com/GauravPandeyLab/KinAce and was also deposited at https://zenodo.org/doi/10.5281/zenodo.10212985 on November 28, 2023.

### Method Detail

#### Web portal construction

The KinAce portal was built as a Shiny^61^ web app in the R ecosystem. The different tabs and their layouts were constructed using the *flexdashboard*^62^ package. Network visualizations were constructed using the *visNetwork*^63^ package and layouts are computed using the *igraph*^64^ package.

The portal is deployed on Amazon Web services under the KinaMetrix domain (https://kinace.kinametrix.com/).

### Quantification and Statistical Analysis

#### Functional analyses of kinase substrates

To examine kinase-modulated cellular functions represented in the KinAce dataset, we performed gene set enrichment on all substrates included against InterPro domains^38^ and KEGG pathways^28^. Specifically, we used the GSEAPy^65^ software to run these enrichment analyses on the Enrichr^56^ platform against the InterPro_Domains_2019 and KEGG_2021_Human gene set libraries. We used the hypergeometric test at a significance level of 0.05 with correction for multiple hypotheses testing applied using the Benjamini-Hochberg procedure^66^.

### Key Resources

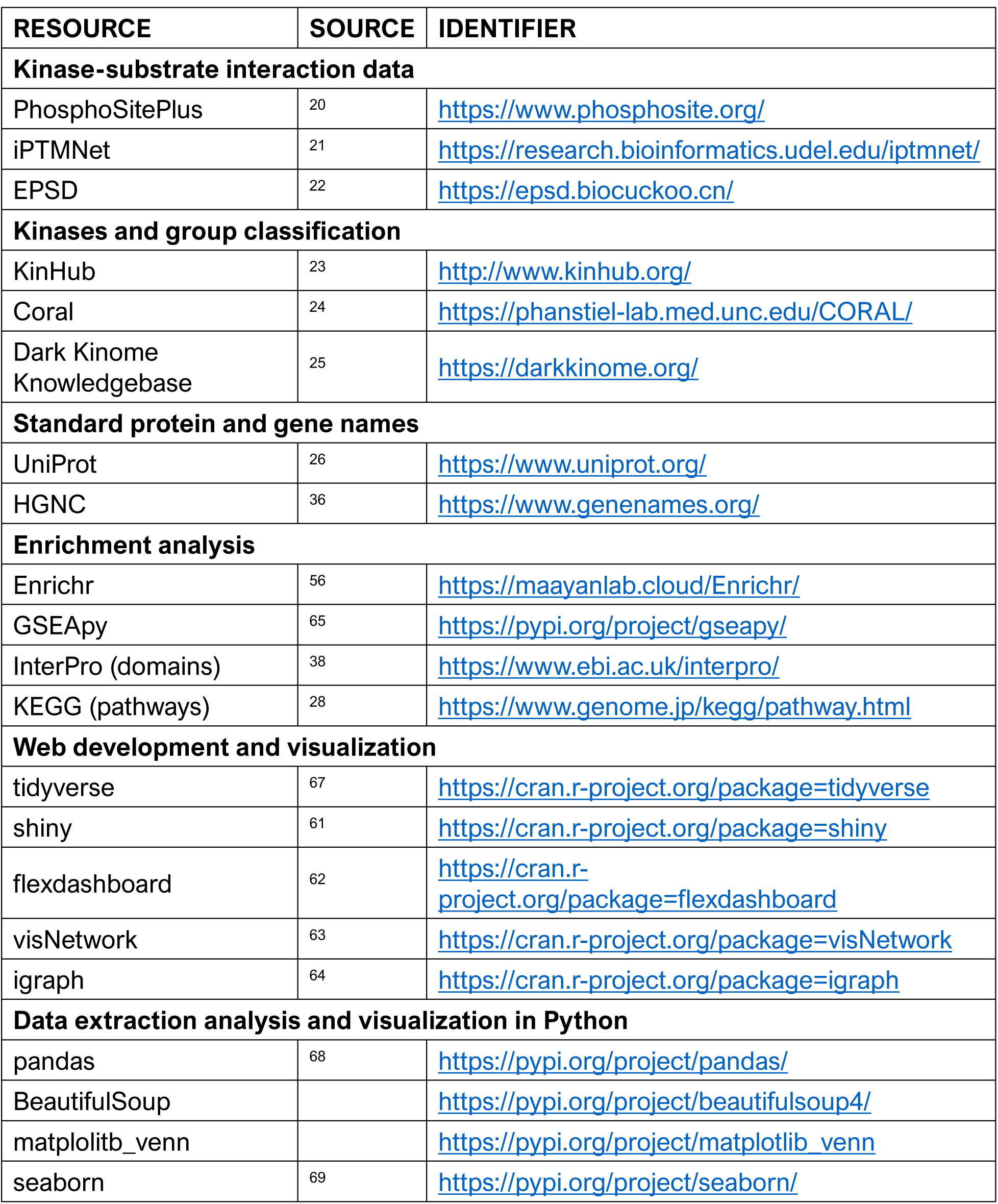

## Supplementary Figures

**Figure S1:**
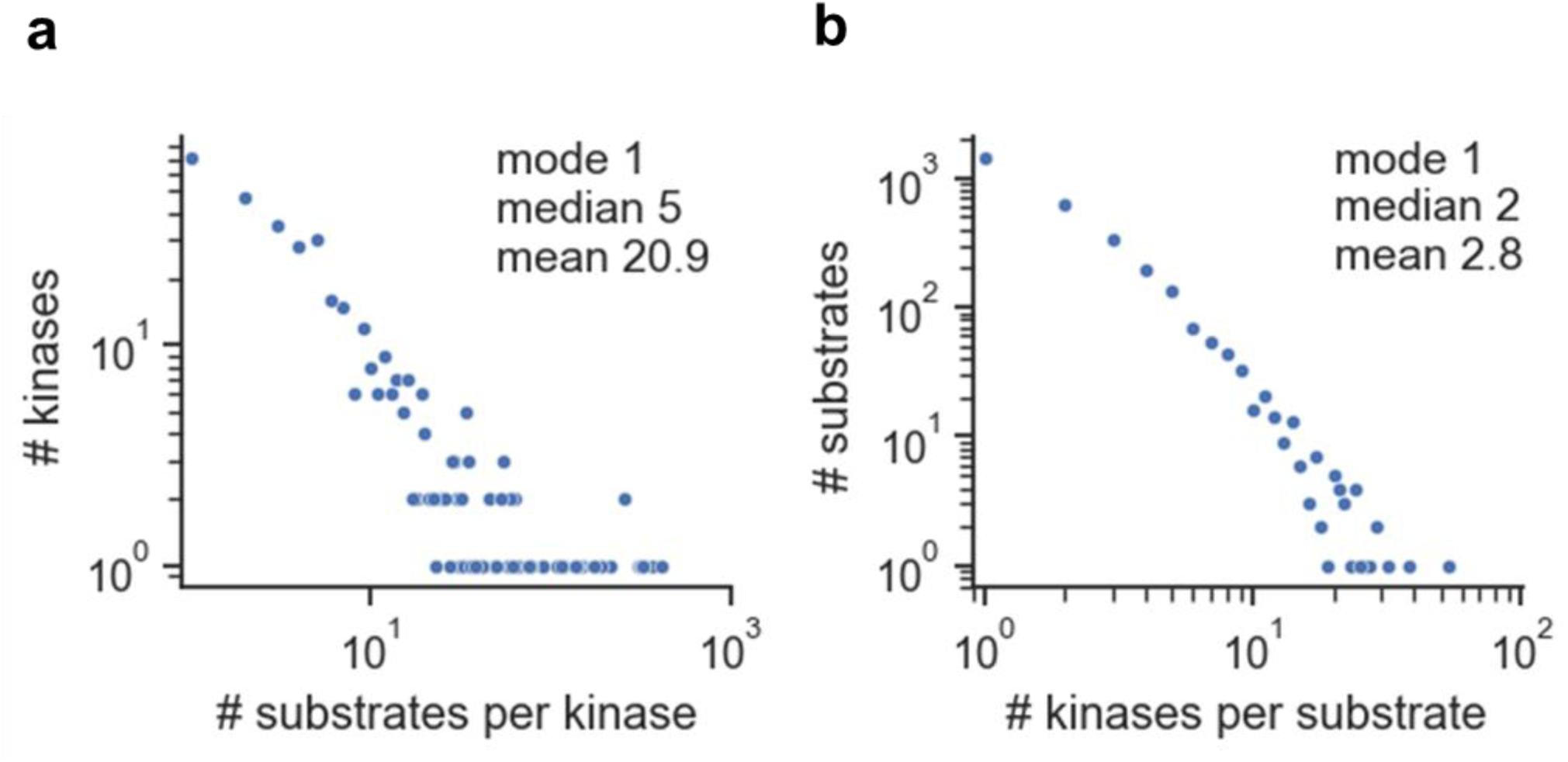
Degree distributions in the KinAce dataset. **(a) Substrates per kinase**. **(b) Kinases per substrate**. Both distributions have heavy right tails, suggesting that a small number of kinases and substrates dominate the set of interactions.

**Figure S2:**
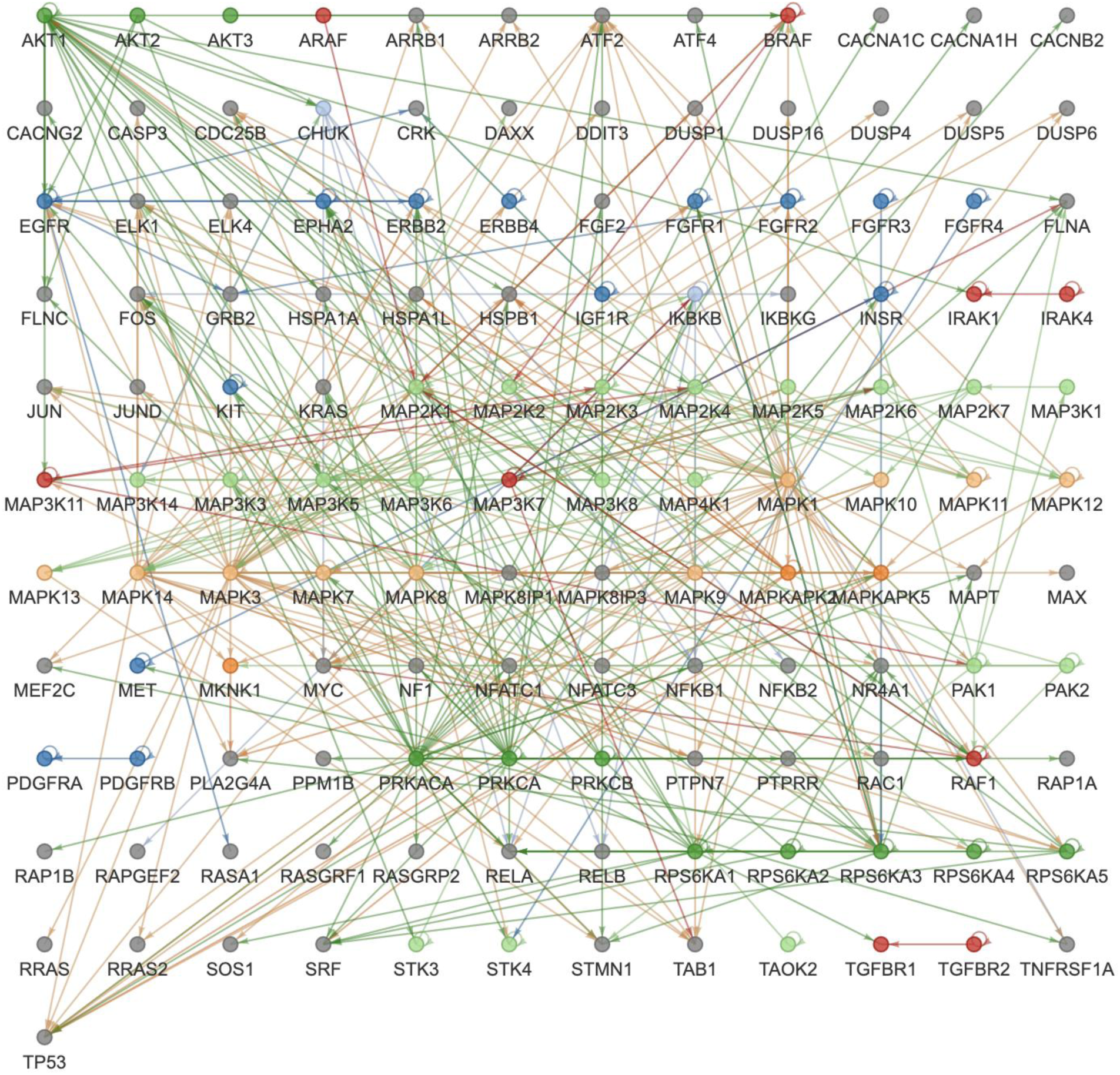
Visualization of the MAPK signaling pathway. The network shown when MAPK signaling is selected in the Pathways tab and grid layout is applied.

**Figure S3:**
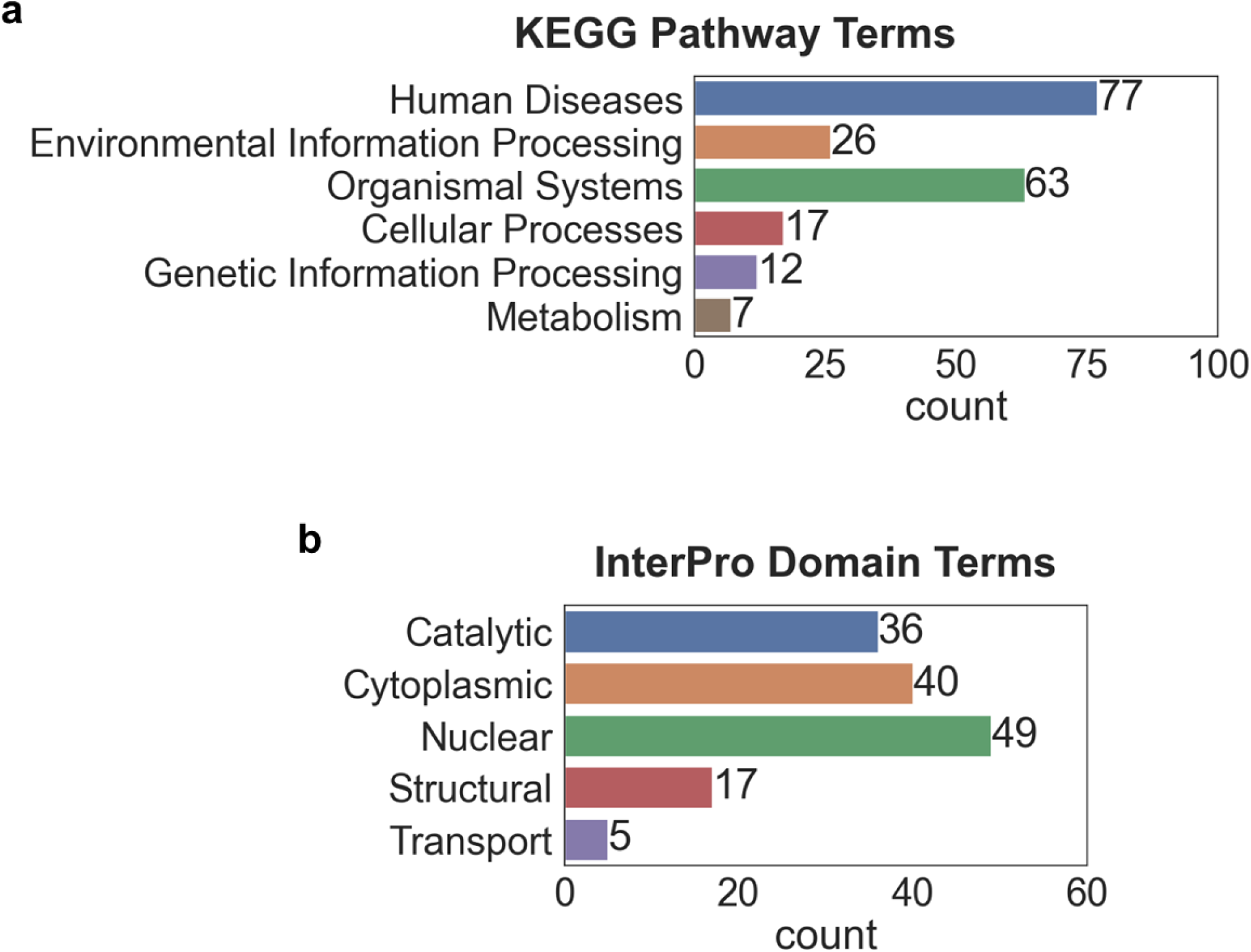
Enrichment analyses of kinase substrates in the KinAce dataset. **(a) KEGG pathways.** This analysis highlighted 201 enriched pathways in six KEGG categories, shown here. The full set of enriched pathways and their p-values are provided in **Appendix 1. (b) InterPro domains.** This analysis highlighted 145 enriched domain terms in five functional categories, shown here. The full set of enriched domain terms and their p-values are provided in Appendix 2.

**Figure S4:**
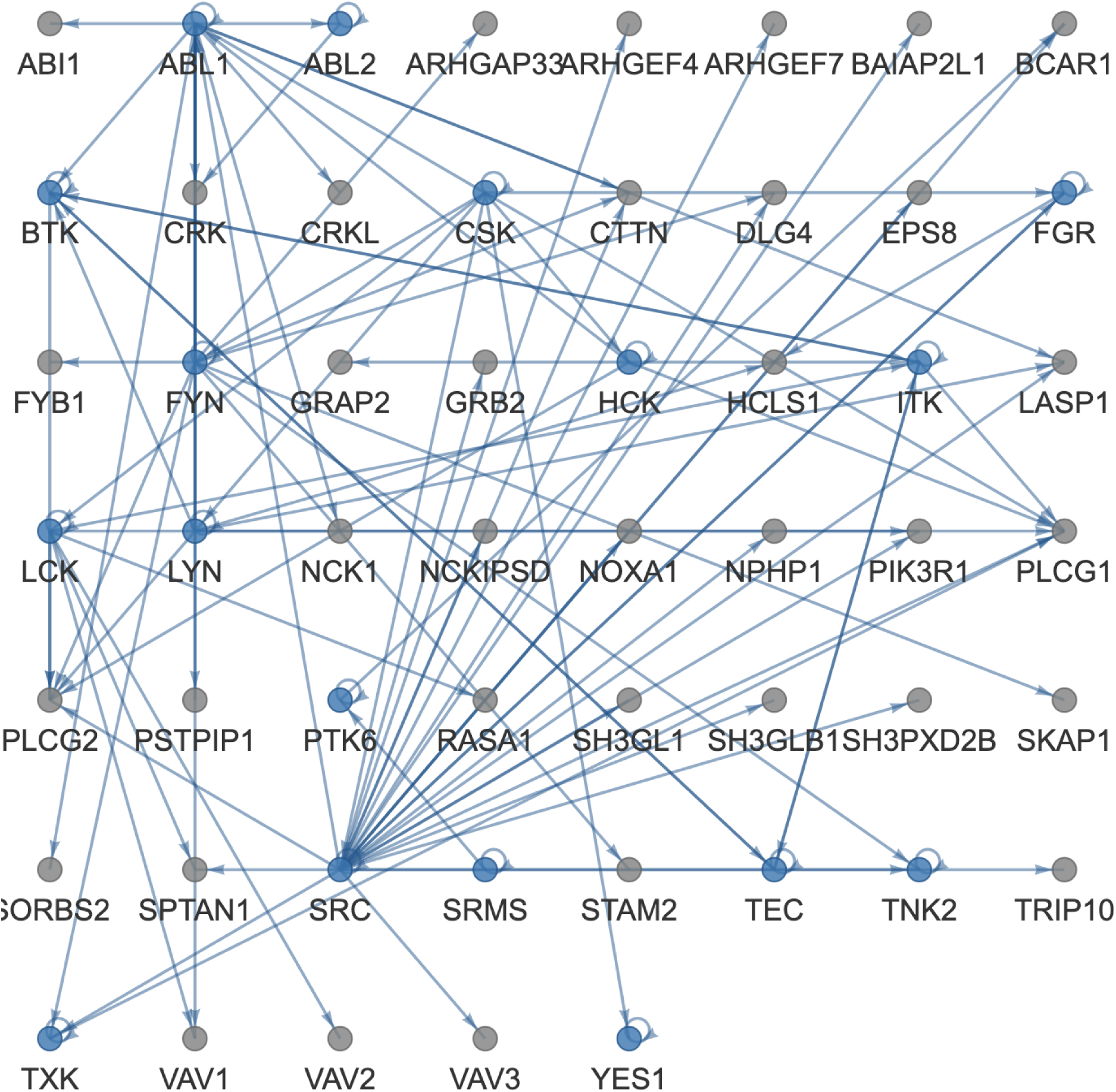
Visualization of proteins with SH3 domain. The network shown when SH3 domain is selected in the Domains tab and grid layout is applied. Note the prevalence of Src-type kinases, such as SRC, FYN, FGR, LCK and LYN.

## Appendix 1: Results of pathway enrichment analysis of kinase substrates in the KinAce dataset

| KEGG Term | KEGG Category | Adjusted p-value |
| --- | --- | --- |
| Pathways in cancer | Human Diseases | 1.53E-70 |
| MAPK signaling pathway | Environmental Information Processing | 6.87E-61 |
| Hepatitis B | Human Diseases | 1.39E-54 |
| Neurotrophin signaling pathway | Organismal Systems | 3.73E-53 |
| MicroRNAs in cancer | Human Diseases | 5.47E-53 |
| Human T-cell leukemia virus 1 infection | Human Diseases | 1.97E-49 |
| Shigellosis | Human Diseases | 3.75E-49 |
| Yersinia infection | Human Diseases | 3.68E-48 |
| Viral carcinogenesis | Human Diseases | 1.22E-47 |
| Proteoglycans in cancer | Human Diseases | 6.21E-44 |
| Cell cycle | Cellular Processes | 1.69E-43 |
| T cell receptor signaling pathway | Organismal Systems | 2.72E-43 |
| Lipid and atherosclerosis | Human Diseases | 7.22E-43 |
| Cellular senescence | Cellular Processes | 2.87E-42 |
| ErbB signaling pathway | Environmental Information Processing | 1.63E-41 |
| Insulin signaling pathway | Organismal Systems | 2.34E-41 |
| Pathways of neurodegeneration | Human Diseases | 2.27E-39 |
| FoxO signaling pathway | Environmental Information Processing | 2.53E-39 |
| Epstein-Barr virus infection | Human Diseases | 1.30E-38 |
| Chronic myeloid leukemia | Human Diseases | 1.72E-38 |
| Focal adhesion | Cellular Processes | 4.44E-38 |
| PI3K-Akt signaling pathway | Environmental Information Processing | 5.12E-38 |
| Human immunodeficiency virus 1 infection | Human Diseases | 1.11E-37 |
| Kaposi sarcoma-associated herpesvirus infection | Human Diseases | 3.11E-37 |
| Salmonella infection | Human Diseases | 3.68E-37 |
| PD-L1 expression and PD-1 checkpoint pathway in cancer | Human Diseases | 3.98E-36 |
| Osteoclast differentiation | Organismal Systems | 4.24E-36 |
| Autophagy | Cellular Processes | 7.90E-36 |
| Regulation of actin cytoskeleton | Cellular Processes | 1.89E-35 |
| Measles | Human Diseases | 3.70E-35 |
| Hepatitis C | Human Diseases | 5.04E-34 |
| Apoptosis | Cellular Processes | 7.65E-34 |
| Ras signaling pathway | Environmental Information Processing | 9.02E-34 |
| Prostate cancer | Human Diseases | 1.63E-33 |
| Pancreatic cancer | Human Diseases | 2.84E-33 |
| Human cytomegalovirus infection | Human Diseases | 6.36E-32 |
| Human papillomavirus infection | Human Diseases | 4.22E-31 |
| Alzheimer disease | Human Diseases | 8.93E-31 |
| Colorectal cancer | Human Diseases | 1.27E-30 |
| cAMP signaling pathway | Environmental Information Processing | 3.14E-30 |
| Insulin resistance | Human Diseases | 3.60E-30 |
| Rap1 signaling pathway | Environmental Information Processing | 4.43E-30 |
| Fc gamma R-mediated phagocytosis | Organismal Systems | 4.60E-30 |
| Pathogenic Escherichia coli infection | Human Diseases | 1.05E-29 |
| Growth hormone synthesis, secretion and action | Organismal Systems | 2.75E-29 |
| Longevity regulating pathway | Organismal Systems | 2.12E-28 |
| Acute myeloid leukemia | Human Diseases | 3.31E-28 |
| C-type lectin receptor signaling pathway | Organismal Systems | 1.07E-27 |
| Renal cell carcinoma | Human Diseases | 1.09E-27 |
| AGE-RAGE signaling pathway in diabetic complications | Human Diseases | 3.65E-27 |
| AMPK signaling pathway | Environmental Information Processing | 3.69E-27 |
| HIF-1 signaling pathway | Environmental Information Processing | 5.58E-27 |
| Non-small cell lung cancer | Human Diseases | 6.44E-27 |
| Chemokine signaling pathway | Organismal Systems | 1.02E-26 |
| Sphingolipid signaling pathway | Environmental Information Processing | 1.36E-26 |
| Tight junction | Cellular Processes | 4.33E-26 |
| Thyroid hormone signaling pathway | Organismal Systems | 5.27E-26 |
| Transcriptional misregulation in cancer | Human Diseases | 1.02E-25 |
| Endocytosis | Cellular Processes | 1.09E-25 |
| Prolactin signaling pathway | Organismal Systems | 1.32E-25 |
| Central carbon metabolism in cancer | Human Diseases | 1.32E-25 |
| Glioma | Human Diseases | 1.32E-25 |
| mTOR signaling pathway | Environmental Information Processing | 1.42E-25 |
| Adherens junction | Cellular Processes | 3.69E-25 |
| Leukocyte transendothelial migration | Organismal Systems | 6.92E-25 |
| Endometrial cancer | Human Diseases | 1.93E-24 |
| TNF signaling pathway | Environmental Information Processing | 2.63E-24 |
| Hepatocellular carcinoma | Human Diseases | 3.89E-24 |
| Axon guidance | Organismal Systems | 4.47E-24 |
| Fc epsilon RI signaling pathway | Organismal Systems | 4.55E-24 |
| Fluid shear stress and atherosclerosis | Human Diseases | 5.30E-24 |
| NOD-like receptor signaling pathway | Organismal Systems | 1.38E-23 |
| Oxytocin signaling pathway | Organismal Systems | 1.50E-23 |
| Breast cancer | Human Diseases | 2.06E-23 |
| Mitophagy | Cellular Processes | 3.38E-23 |
| Neutrophil extracellular trap formation | Organismal Systems | 5.23E-23 |
| Platelet activation | Organismal Systems | 5.32E-23 |
| cGMP-PKG signaling pathway | Environmental Information Processing | 6.04E-23 |
| B cell receptor signaling pathway | Organismal Systems | 1.21E-22 |
| Th17 cell differentiation | Organismal Systems | 2.52E-22 |
| Progesterone-mediated oocyte maturation | Organismal Systems | 9.80E-22 |
| Gastric cancer | Human Diseases | 1.03E-21 |
| Apelin signaling pathway | Environmental Information Processing | 2.06E-21 |
| Chemical carcinogenesis | Human Diseases | 4.04E-21 |
| Toxoplasmosis | Human Diseases | 5.61E-21 |
| Bacterial invasion of epithelial cells | Human Diseases | 8.79E-21 |
| Necroptosis | Cellular Processes | 9.11E-21 |
| Toll-like receptor signaling pathway | Organismal Systems | 1.25E-20 |
| Phospholipase D signaling pathway | Environmental Information Processing | 5.50E-20 |
| Spinocerebellar ataxia | Human Diseases | 6.71E-20 |
| GnRH signaling pathway | Organismal Systems | 1.00E-19 |
| Hippo signaling pathway | Environmental Information Processing | 1.33E-19 |
| NF-kappa B signaling pathway | Environmental Information Processing | 1.63E-19 |
| Long-term potentiation | Organismal Systems | 1.85E-19 |
| Signaling pathways regulating pluripotency of stem cells | Cellular Processes | 2.06E-19 |
| Amyotrophic lateral sclerosis | Human Diseases | 2.67E-19 |
| Influenza A | Human Diseases | 2.74E-19 |
| Choline metabolism in cancer | Human Diseases | 3.29E-19 |
| Small cell lung cancer | Human Diseases | 6.81E-19 |
| VEGF signaling pathway | Environmental Information Processing | 7.79E-19 |
| Parathyroid hormone synthesis, secretion and action | Organismal Systems | 8.97E-19 |
| Diabetic cardiomyopathy | Human Diseases | 1.40E-18 |
| Oocyte meiosis | Cellular Processes | 1.42E-18 |
| p53 signaling pathway | Cellular Processes | 2.22E-18 |
| Cushing syndrome | Human Diseases | 2.79E-18 |
| Alcoholism | Human Diseases | 1.34E-17 |
| Bladder cancer | Human Diseases | 1.53E-17 |
| Chagas disease | Human Diseases | 3.69E-17 |
| Non-alcoholic fatty liver disease | Human Diseases | 4.86E-17 |
| Estrogen signaling pathway | Organismal Systems | 5.64E-17 |
| Adipocytokine signaling pathway | Organismal Systems | 6.68E-17 |
| IL-17 signaling pathway | Organismal Systems | 9.29E-17 |
| Tuberculosis | Human Diseases | 9.95E-17 |
| Calcium signaling pathway | Environmental Information Processing | 1.29E-16 |
| Relaxin signaling pathway | Organismal Systems | 1.74E-16 |
| Melanoma | Human Diseases | 5.32E-16 |
| Th1 and Th2 cell differentiation | Organismal Systems | 1.10E-15 |
| Coronavirus disease | Human Diseases | 1.55E-15 |
| Adrenergic signaling in cardiomyocytes | Organismal Systems | 1.70E-15 |
| Huntington disease | Human Diseases | 9.58E-15 |
| Parkinson disease | Human Diseases | 1.99E-14 |
| Type II diabetes mellitus | Human Diseases | 3.18E-14 |
| Prion disease | Human Diseases | 4.12E-14 |
| Leishmaniasis | Human Diseases | 4.34E-14 |
| Thyroid cancer | Human Diseases | 1.21E-13 |
| RIG-I-like receptor signaling pathway | Organismal Systems | 1.88E-13 |
| Glucagon signaling pathway | Organismal Systems | 1.93E-13 |
| Pertussis | Human Diseases | 2.64E-13 |
| Inflammatory mediator regulation of TRP channels | Organismal Systems | 3.86E-13 |
| GnRH secretion | Organismal Systems | 3.93E-13 |
| Dopaminergic synapse | Organismal Systems | 4.34E-13 |
| Wnt signaling pathway | Environmental Information Processing | 4.54E-13 |
| JAK-STAT signaling pathway | Environmental Information Processing | 2.21E-12 |
| Gap junction | Cellular Processes | 3.70E-12 |
| Ubiquitin mediated proteolysis | Genetic Information Processing | 3.86E-12 |
| Long-term depression | Organismal Systems | 5.55E-12 |
| Cholinergic synapse | Organismal Systems | 1.42E-11 |
| Natural killer cell mediated cytotoxicity | Organismal Systems | 1.81E-11 |
| Amphetamine addiction | Human Diseases | 3.98E-11 |
| Aldosterone synthesis and secretion | Organismal Systems | 2.44E-10 |
| Homologous recombination | Genetic Information Processing | 2.67E-09 |
| Non-homologous end-joining | Genetic Information Processing | 2.95E-09 |
| Vascular smooth muscle contraction | Organismal Systems | 6.88E-09 |
| Cocaine addiction | Human Diseases | 7.13E-09 |
| Epithelial cell signaling in *Helicobacter pylori* infection | Human Diseases | 8.19E-09 |
| Circadian entrainment | Organismal Systems | 1.34E-07 |
| RNA transport | Genetic Information Processing | 1.44E-07 |
| Aldosterone-regulated sodium reabsorption | Organismal Systems | 2.66E-07 |
| Melanogenesis | Organismal Systems | 4.29E-07 |
| Regulation of lipolysis in adipocytes | Organismal Systems | 6.28E-07 |
| Fanconi anemia pathway | Genetic Information Processing | 1.24E-06 |
| Legionellosis | Human Diseases | 1.38E-06 |
| Serotonergic synapse | Organismal Systems | 2.47E-06 |
| Primary immunodeficiency | Human Diseases | 2.63E-06 |
| Notch signaling pathway | Environmental Information Processing | 2.88E-06 |
| Circadian rhythm | Organismal Systems | 7.54E-06 |
| Thermogenesis | Organismal Systems | 1.19E-05 |
| Endocrine and other factor-regulated calcium reabsorption | Organismal Systems | 2.07E-05 |
| Protein processing in endoplasmic reticulum | Genetic Information Processing | 2.19E-05 |
| Phosphatidylinositol signaling system | Environmental Information Processing | 3.55E-05 |
| Hypertrophic cardiomyopathy | Human Diseases | 5.45E-05 |
| Insulin secretion | Organismal Systems | 5.98E-05 |
| Arrhythmogenic right ventricular cardiomyopathy | Human Diseases | 1.63E-04 |
| Cell adhesion molecules | Environmental Information Processing | 1.84E-04 |
| Base excision repair | Genetic Information Processing | 2.45E-04 |
| Gastric acid secretion | Organismal Systems | 3.56E-04 |
| Dilated cardiomyopathy | Human Diseases | 4.06E-04 |
| Hedgehog signaling pathway | Environmental Information Processing | 4.39E-04 |
| Inositol phosphate metabolism | Metabolism | 4.89E-04 |
| Renin secretion | Organismal Systems | 5.43E-04 |
| Retrograde endocannabinoid signaling | Organismal Systems | 5.96E-04 |
| Cortisol synthesis and secretion | Organismal Systems | 5.96E-04 |
| TGF-beta signaling pathway | Environmental Information Processing | 6.58E-04 |
| Vasopressin-regulated water reabsorption | Organismal Systems | 6.68E-04 |
| Glutamatergic synapse | Organismal Systems | 7.52E-04 |
| Glycolysis / Gluconeogenesis | Metabolism | 9.21E-04 |
| Ferroptosis | Cellular Processes | 9.21E-04 |
| Morphine addiction | Human Diseases | 1.07E-03 |
| Viral myocarditis | Human Diseases | 1.13E-03 |
| Salivary secretion | Organismal Systems | 1.51E-03 |
| Spliceosome | Genetic Information Processing | 1.59E-03 |
| Cytosolic DNA-sensing pathway | Organismal Systems | 2.18E-03 |
| Phagosome | Cellular Processes | 2.23E-03 |
| DNA replication | Genetic Information Processing | 2.36E-03 |
| Amoebiasis | Human Diseases | 2.47E-03 |
| Basal cell carcinoma | Human Diseases | 2.64E-03 |
| Purine metabolism | Metabolism | 2.76E-03 |
| mRNA surveillance pathway | Genetic Information Processing | 3.30E-03 |
| Systemic lupus erythematosus | Human Diseases | 3.57E-03 |
| Inflammatory bowel disease | Human Diseases | 3.86E-03 |
| Carbohydrate digestion and absorption | Organismal Systems | 4.17E-03 |
| Pyruvate metabolism | Metabolism | 4.17E-03 |
| Maturity onset diabetes of the young | Human Diseases | 4.63E-03 |
| Mismatch repair | Genetic Information Processing | 6.48E-03 |
| Nucleotide excision repair | Genetic Information Processing | 9.07E-03 |
| Bile secretion | Organismal Systems | 9.17E-03 |
| Pancreatic secretion | Organismal Systems | 1.15E-02 |
| Pentose phosphate pathway | Metabolism | 1.44E-02 |
| Citrate cycle (TCA cycle) | Metabolism | 1.44E-02 |
| Thyroid hormone synthesis | Organismal Systems | 1.95E-02 |
| Hematopoietic cell lineage | Organismal Systems | 4.65E-02 |
| Folate biosynthesis | Metabolism | 4.76E-02 |

## Appendix 2: Results of domain enrichment analysis of kinase substrates in the KinAce dataset

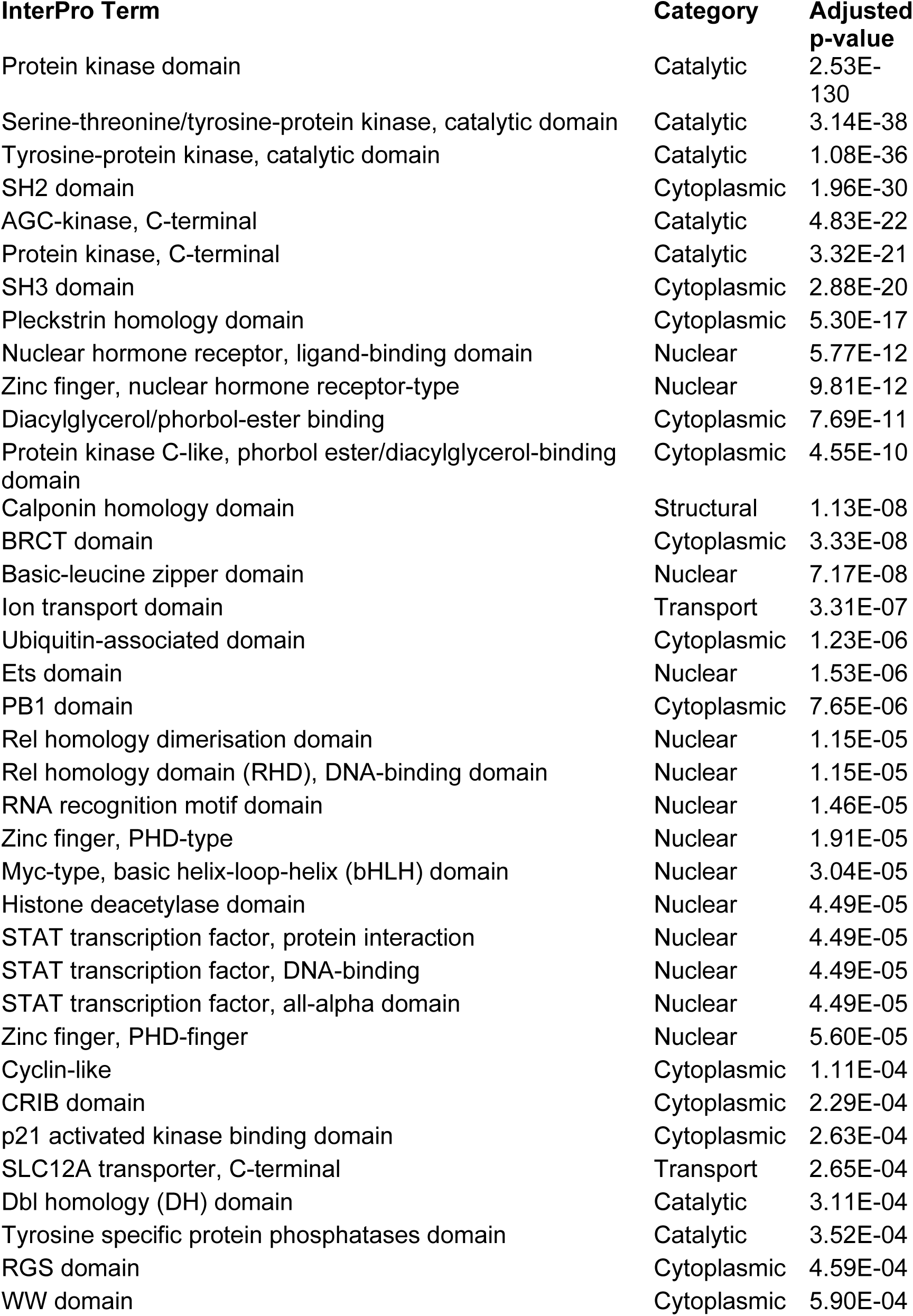
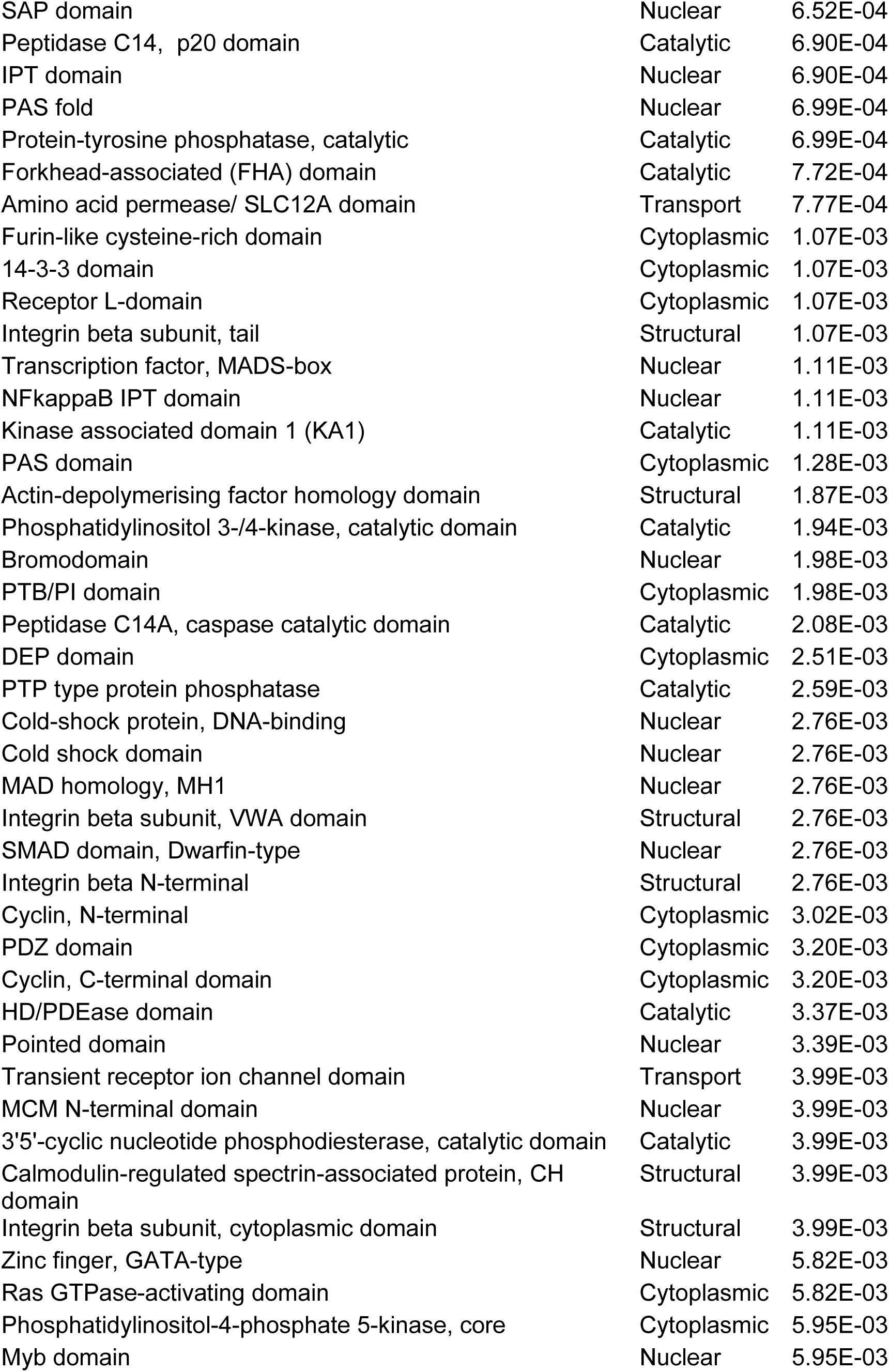
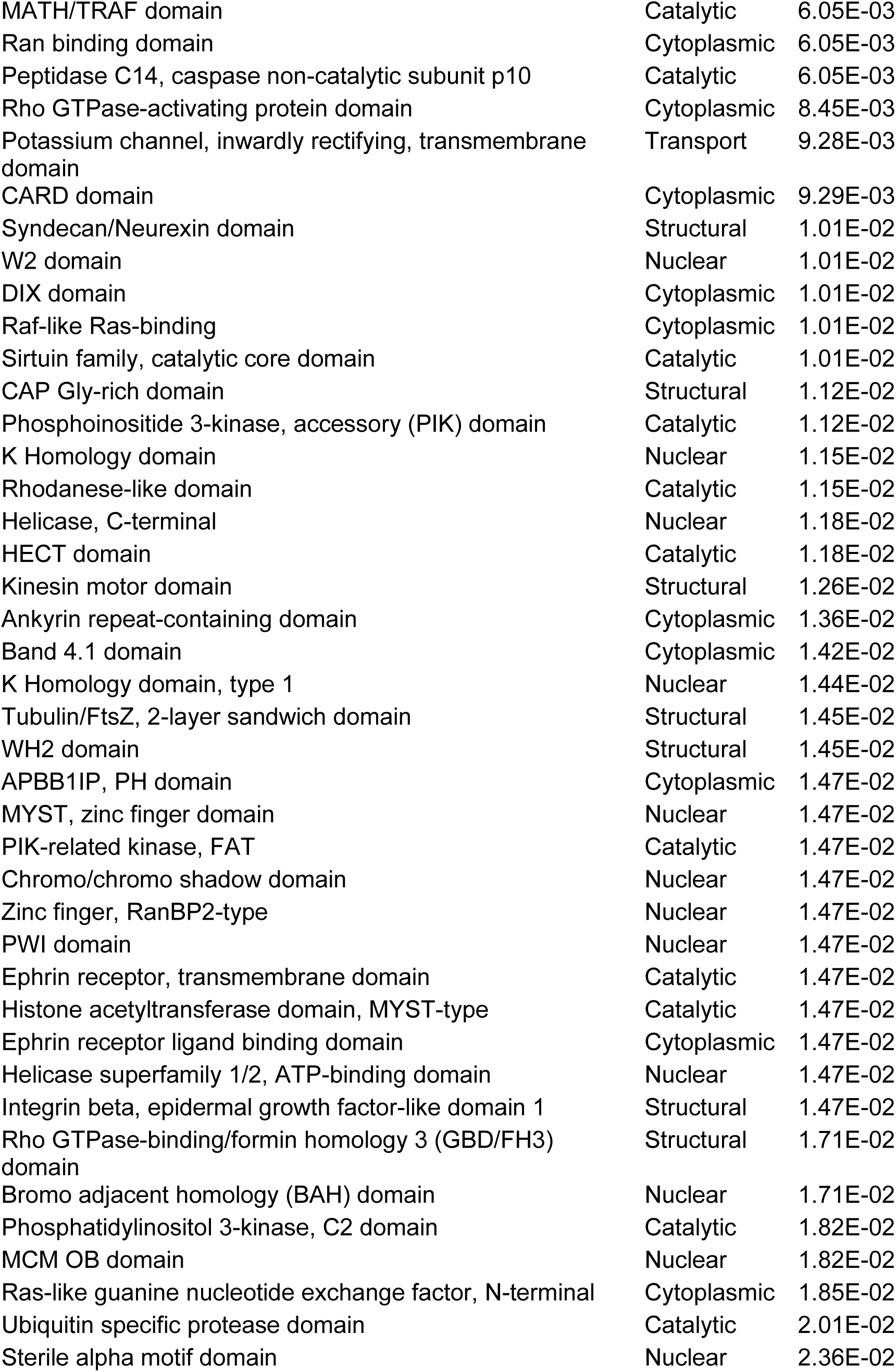
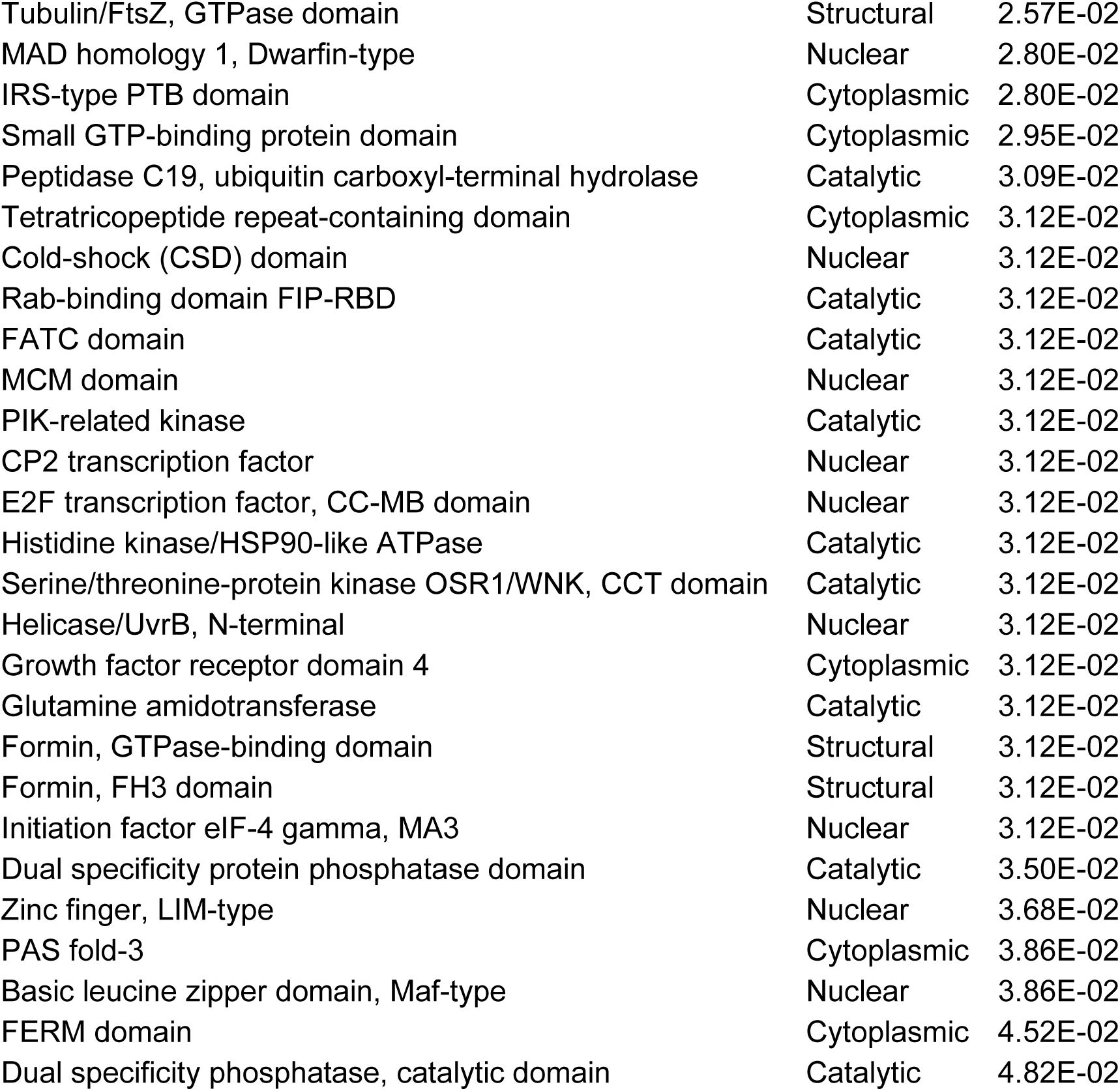

